## Supplemental Materials for "Territoriality modulates the coevolution of cooperative breeding and female song in songbirds"

**Supplementary Information for Snyder, Loughran-Pierce, and Creanza (2025), "Territoriality modulates the coevolution of cooperative breeding and female song in songbirds": Supplemental Tables 1–25, Supplemental Figures 1–2, Supplementary Results Text**

**Supplemental Table 1:** Binary states of each sociality metric. The last six rows provide the cooperative breeding assignments used in this manuscript. The number of species is the number of Oscine species with the noted classification, separated by state. In all analyses where the specific cooperative breeding classification method is not noted (i.e. in the main text), HighConfidence_Coop was used.

| Original source column name | Source | Binarized column name in Supplemental Dataset 2 | Source values assigned state "0" or "1" | State "0" and state "1" categories in this study | Number of oscine species assigned to state |
|---|---|---|---|---|---|
| colonial | Griesser et al (2023) | Griesser2023.Colonial01 | acolonial (0) | Non-colonial | 404 |
| | | | colonial (1) | Colonial | 83 |
| parental.care.mode | Griesser et al (2017) | Griesser2017FamilialLiving | no_fam (0) | Non-familial living | 577 |
| | | | coop_families, family (1) | Familial living | 626 |
| grouping | Griesser et al (2023) | Griesser2023.Asocial0VsSocial1 | asocial (0) | Asocial | 47 |
| | | | pair, small_groups, large_groups (1) | Social | 409 |
| grouping | Griesser et al (2023) | Griesser2023.GroupsLargerThanPair | asocial, pair (0) | Groups pair or smaller | 88 |
| | | | small_groups, large_groups (1) | Groups larger than pair | 368 |
| grouping | Griesser et al (2023) | Griesser2023.LargestGroupSizes | asocial, pair, small_groups (0) | Smaller groups | 306 |
| | | | large_groups (1) | Largest group sizes | 150 |
| social_bonds | Griesser et al (2023) | Griesser2023.LongSocialBonds | a-short, b-season (0) | Short social bonds | 170 |
| | | | c-long (1) | Season or shorter social bond | 317 |
| social_bonds | Griesser et al (2023) | Griesser2023.SeasonOrLongerSocialBonds | a-short (0) | Season or shorter social bonds | 18 |
| | | | b-season, c-long (1) | Long social bonds | 469 |
| caretakers | Griesser et al (2023) | Griesser2023.MoreThanTwoCaretakers | <= 2 (0) | Two or fewer caretakers | 411 |
| | | | > 2 (1) | More than two caretakers | 76 |
| caretakers | Griesser et al (2023) | Griesser2023.TwoOrMoreCaretakers | < 2 (0) | Fewer than two caretakers | 24 |
| | | | >= 2 (1) | Two or more caretakers | 463 |
| Final.polygyny | Snyder and Creanza (2019) | Final.polygyny | 0 (Socially monogamous) | Social monogamy | 626 |
| | | | 1 (Socially polygynous) | Social polygyny | 108 |
| Territory | Tobias et al (2016) | Territory_YearRound | 1, 2 (0) | No territoriality, weak territoriality, or seasonal territoriality | 3186 |
| | | | 3 (1) | Year-round territoriality | 1246 |
| Current study | Current study | TerritorialityWeakVsStrong | See methods | No territory or territory not actively defended | 604 |
| | | | See methods | Territory actively defended by individual, pair, or group | 1560 |
| Current study | Current study | HighConfidence_Coop | See methods | Non-cooperative | 3904 |
| | | | See methods | Cooperative | 469 |
| Current study | Current study | MeanCoopOmitTies | See methods | Non-cooperative | 3940 |
| | | | See methods | Cooperative | 493 |
| Current study | Current study | MeanCoopTie2Coop | See methods | Non-cooperative | 3940 |
| | | | See methods | Cooperative | 554 |
| Current study | Current study | MeanCoopTie2Noncoop | See methods | Non-cooperative | 4001 |
| | | | See methods | Cooperative | 493 |
| Current study | Current study | AnyCoopEqualsCoop | See methods | Non-cooperative | 3859 |
| | | | See methods | Cooperative | 635 |
| Current study | Current study | AnyNoncoopEqualsNoncoop | See methods | Non-cooperative | 4060 |
| | | | See methods | Cooperative | 434 |

**Supplemental Table 2:** Species counts for each combination of female song presence and breeding system and sociality metrics. See Supplemental Table 1 for sources.

| Sociality variables | Female Song Absent | Female Song Present | Non-cooperative Breeding | Cooperative Breeding |
|---|---|---|---|---|
| Non-colonial (breeds singly) | 58 | 165 | 328 | 61 |
| Colonial (breed in colonies) | 12 | 35 | 74 | 9 |
| Asocial (individuals usually on their own) | 8 | 26 | 46 | 0 |
| Social (individuals usually with at least one other individual) | 56 | 165 | 325 | 70 |
| Groups pair or smaller (usually on their own or in pairs) | 15 | 49 | 85 | 0 |
| Groups larger than pair (usually in groups of 3 or more) | 49 | 142 | 286 | 70 |
| Smaller groups (individuals usually in groups of 0-30 individuals) | 34 | 139 | 230 | 65 |
| Largest group sizes (individuals usually in groups of >30 individuals) | 30 | 52 | 141 | 5 |
| Short social bonds (bonds lasting less than one season) | 11 | 2 | 18 | 0 |
| Season or longer social bonds (bonds lasting one season or longer) | 59 | 198 | 384 | 70 |
| Season or shorter social bonds (bonds lasting at most one season) | 34 | 61 | 161 | 3 |
| Long social bonds (bonds lasting longer than one season) | 36 | 139 | 241 | 67 |
| Two or fewer caretakers (species exhibited mound-nesting (zero caretakers), uniparental care (one caretaker), or biparental care (two caretakers) | 66 | 172 | 401 | 1 |
| More than two caretakers (species had cooperative breeding with more than two caretakers) | 4 | 28 | 1 | 69 |
| Fewer than two caretakers (species exhibited mound-nesting (zero caretakers) or uniparental care (one caretaker)) | 12 | 6 | 24 | 0 |
| Two or more caretakers (species had biparental care or cooperative breeding with more than two caretakers) | 58 | 194 | 378 | 70 |
| Non-familial living (parent–offspring associations do not extend beyond nutritional independence [includes non-kin cooperatively breeding species]) | 103 | 159 | 554 | 11 |
| Familial living (offspring remain with their parents beyond nutritional independence [includes kin-based cooperatively breeding species]) | 59 | 147 | 345 | 249 |
| Social monogamy (<5% of studied males exhibited polygyny) | 102 | 166 | 531 | 72 |
| Social polygyny (≥5% of males exhibited polygyny) | 32 | 23 | 104 | 2 |
| Asocial (usually on their own) | 8 | 26 | 46 | 0 |
| Pair (usually in pairs) | 7 | 23 | 39 | 0 |
| Small groups (usually 3 to 30 individuals (approximating personalized groups where group members know each other individually, including family groups) | 19 | 90 | 145 | 65 |
| Large groups (usually more than 30 individuals; more ephemeral or anonymous groups) | 30 | 52 | 141 | 5 |
| Weak, seasonal, or no territoriality | 349 | 313 | 2807 | 218 |
| Year-round territoriality | 47 | 382 | 881 | 237 |
| Weak or no territoriality | 106 | 71 | 543 | 49 |
| Active defense of territory (seasonally or year-round) | 201 | 541 | 1162 | 258 |

**Supplemental Table 3:** Results of phylANOVA testing whether song features differ between states of different sociality traits. Continuous song traits were natural-log-transformed before analysis. Each trait pair (a combination of a discrete social trait and a continuous song trait) was tested using phytools::phylANOVA() for 50,000 simulations each. All song traits were tested relative to our main cooperative breeding dataset (HighConfidence_Coop, denoted in pink), but only song repertoire size was tested with other classification methods (yellow), with other social traits (blue), or with data obtained from individual cooperative breeding sources (purple) since none of the other song traits (syllable repertoire size, syllables per song, song duration, inter-song interval) showed clear trends relative to cooperative breeding in either phylANOVA or Brownie.

| Social Trait | # of states of social trait | Song Trait | Number of species with data for both the social and song traits | phylANOVA p-value |
|---|---|---|---|---|
| HighConfidence_Coop | 2 | Syllable.rep.final | 118 | 0.34825 |
| HighConfidence_Coop | 2 | Song.rep.final | 209 | 0.9772 |
| HighConfidence_Coop | 2 | Syll.song.final | 163 | 0.82805 |
| HighConfidence_Coop | 2 | Duration.final | 218 | 0.35155 |
| HighConfidence_Coop | 2 | Interval.final | 123 | 0.9097 |
| MeanCoopTie2Noncoop | 2 | Song.rep.final | 217 | 0.96575 |
| MeanCoopTie2Coop | 2 | Song.rep.final | 217 | 0.45575 |
| MeanCoopOmitTies | 2 | Song.rep.final | 214 | 0.9358 |
| AnyCoopEqualsCoop | 2 | Song.rep.final | 217 | 0.7963 |
| AnyNoncoopEqualsNoncoop | 2 | Song.rep.final | 217 | 0.77096 |
| social_system_Griesser2017 | 3 | Song.rep.final | 156 | 0.93704 |
| social_system_incl_nk_coop_Griesser2017 | 4 | Song.rep.final | 156 | 0.66026 |
| grouping | 4 | Song.rep.final | 127 | 0.72988 |
| Griesser2023.Colonial01 | 2 | Song.rep.final | 129 | 0.79964 |
| Griesser2023.MoreThanTwoCaretakers | 2 | Song.rep.final | 129 | 0.85682 |
| Griesser2023.LongSocialBonds | 2 | Song.rep.final | 129 | 0.65492 |
| Griesser2023.GroupsLargerThanPair | 2 | Song.rep.final | 127 | 0.3345 |
| Griesser2023.TwoOrMoreCaretakers | 2 | Song.rep.final | 129 | 0.67844 |
| Griesser2023.Asocial0vsSocial1 | 2 | Song.rep.final | 127 | 0.26796 |
| Griesser2023.LargestGroupSizes | 2 | Song.rep.final | 127 | 0.56416 |
| Griesser2023.SeasonOrLongerSocialBonds | 2 | Song.rep.final | 129 | 0.64316 |
| Griesser2017FamilialLiving | 2 | Song.rep.final | 156 | 0.78656 |
| Territory (Tobias et al 2016) | 3 | Song.rep.final | 216 | 0.58884 |
| Territory_YearRound | 2 | Song.rep.final | 216 | 0.4243 |
| TerritorialityWeakVsStrong | 2 | Song.rep.final | 159 | 0.53198 |
| Cooperative breeding classifications from Biagolini et al (2017) | 2 | Song.rep.final | 73 | 0.8115 |
| Cooperative breeding classifications from Downing et al (2015) | 2 | Song.rep.final | 71 | 0.3987 |
| Cooperative breeding classifications from Cockburn (2016) | 2 | Song.rep.final | 181 | 0.43534 |
| Cooperative breeding classifications from Griesser et al (2017) | 2 | Song.rep.final | 156 | 0.53256 |
| Cooperative breeding classifications from Dale (2015) | 2 | Song.rep.final | 199 | 0.61664 |
| Cooperative breeding classifications from Cornwallis (2017) | 2 | Song.rep.final | 204 | 0.6146 |
| Cooperative breeding classifications from Jetz & Rubenstein (2011) | 2 | Song.rep.final | 215 | 0.2319 |

**Supplemental Table 4:** Results assessing whether rates of song-feature evolution differ based on the predicted ancestral state of multiple binary sociality metrics (for multi-state metrics, see Supplemental Table 6). For each combination of song feature and sociality metric, out of 500 simulations per combination run using the Brownie algorithm (brownie.lite, R package: phytools), the Higher Rate State is determined as the discrete state in which the continuous trait evolved faster in at least 50% of the simulations, and the Lower Rate State is the other state. Cells highlighted in green indicate that at least 95% of simulations followed that Higher/Lower rate trend. The overall *p*-value was obtained by taking the mean log-likelihood from all of the Equal Rates simulations and the mean log-likelihood from all of the All Rates Different simulations and performing a likelihood ratio test (*p*-values <0.05 highlighted in yellow). Results are provided for cooperative breeding system for all song features (pink). Song repertoire, syllable repertoire, and syllables per song each have at least one metric that is sufficiently significant, so we repeated the analysis using the minimum and maximum values per species for each; only song repertoire size had consistently significant results using min/max values, as well as across 500 iterations of resampled min/median/max values (orange), so we can only confidently conclude that song repertoire size appears to evolve at a significantly higher rate in non-cooperative breeding lineages than in cooperatively breeding lineages. We then also tested for state-dependent rates of song repertoire size evolution using alternative methods of cooperative breeding classification (yellow), our alternative consensus phylogeny (light green), multiple unique tree topographies (200 unique trees with the Hackett backbone from BirdTree.org, 20 simulations per tree; light green), other measures of sociality from Griesser et al (2017) and Griesser et al (2023) (see Supplemental Table 1; blue), and cooperative breeding data from individual sources (purple). The names of the social traits and song traits correspond to column names in Supplemental Datasets 1 and 2. Species counts are in Supplemental Table 3.

| Binary Social Trait | Continuous Song Trait | Higher Rate State | Lower Rate State | Fraction of Sims Following Trend | Brownie *p*-value |
|---|---|---|---|---|---|
| HighConfidence_Coop | Syllable.rep.final | Non-Cooperative | Cooperative | 0.982 | 0.136 |
| HighConfidence_Coop | Song.rep.final | Non-Cooperative | Cooperative | 0.994 | 0.024 |
| HighConfidence_Coop | Syll.song.final | Cooperative | Non-Cooperative | 0.906 | 0 |
| HighConfidence_Coop | Duration.final | Cooperative | Non-Cooperative | 0.704 | 0.0887 |
| HighConfidence_Coop | Interval.final | Non-Cooperative | Cooperative | 0.884 | 0.3797 |
| HighConfidence_Coop | Syllable.rep.min | Non-Cooperative | Cooperative | 0.988 | 0.1447 |
| HighConfidence_Coop | Syllable.rep.max | Non-Cooperative | Cooperative | 0.808 | 0.2336 |
| HighConfidence_Coop | Song.rep.min | Non-Cooperative | Cooperative | 0.992 | 0.026 |
| HighConfidence_Coop | Song.rep.max | Non-Cooperative | Cooperative | 0.994 | 0.028 |
| HighConfidence_Coop | Syll.song.min | Cooperative | Non-Cooperative | 0.696 | 0.2536 |
| HighConfidence_Coop | Syll.song.max | Cooperative | Non-Cooperative | 0.934 | 0 |
| HighConfidence_Coop | Song repertoire (resampled min/median/max values) | Non-Cooperative | Cooperative | 0.993 | 0.0332 |
| MeanCoopTie2Noncoop | Song.rep.final | Non-Cooperative | Cooperative | 0.992 | 0.0062 |
| MeanCoopTie2Coop | Song.rep.final | Non-Cooperative | Cooperative | 1 | 0.0029 |
| MeanCoopOmitTies | Song.rep.final | Non-Cooperative | Cooperative | 0.994 | 0.0077 |
| AnyCoopEqualsCoop | Song.rep.final | Non-Cooperative | Cooperative | 0.89 | 0.2222 |
| AnyNoncoopEqualsNoncoop | Song.rep.final | Non-Cooperative | Cooperative | 0.984 | 0.0194 |
| HighConfidence_Coop (pooled results across 200 randomly sampled trees) | Song.rep.final | Non-Cooperative | Cooperative | 0.98883333 | 0.032 |
| HighConfidence_Coop (using alternative consensus tree) | Song.rep.final | Non-Cooperative | Cooperative | 0.99 | 0.03 |
| Griesser2023.Asocial0VsSocial1 | Song.rep.final | Pair or group sociality | Asocial | 1 | 0.0723 |
| Griesser2023.GroupsLargerThanPair | Song.rep.final | Asocial or pair | Small or large groups | 1 | 0.171 |
| Griesser2023.LargestGroupSizes | Song.rep.final | Large groups | Asocial, pair, or small groups | 0.53 | 0.6477 |
| Griesser2023.SeasonOrLongerSocialBonds | Song.rep.final | Bonds last less than one season | Season or longer social bonds | 0.89 | 0.3071 |
| Griesser2023.LongSocialBonds | Song.rep.final | Bonds last one season or less | Multi-year bonds | 1 | 0.0079 |
| Griesser2017FamilialLiving | Song.rep.final | Non-Familial Living | Familial Living | 0.998 | 0.0042 |

| | | | | | |
|---|---|---|---|---|---|
| Griesser2023.Colonial01 | Song.rep.final | Colonial | Non-Colonial | 0.99 | 0.115 |
| Griesser2023.MoreThanTwoCaretakers | Song.rep.final | Two or fewer caretakers | More than two caretakers | 0.998 | 0.057 |
| Territory_YearRound | Song.rep.final | Weak, seasonal, or no territoriality | Year-round territoriality | 1 | 0.0106 |
| TerritorialityWeakVsStrong | Song.rep.final | Weak or no territoriality | Strong territoriality | 0.988 | 0.2165 |
| Cooperative breeding classifications from Biagolini et al 2017 (transition rates based on source dataset) | Song.rep.final | Non-Cooperative | Cooperative | 0.996 | 0.092 |
| Cooperative breeding classifications from Biagolini et al 2017 (transition rates based on current study dataset) | Song.rep.final | Non-Cooperative | Cooperative | 0.998 | 0.039 |
| Cooperative breeding classifications from Downing et al (2015) (transition rates based on source dataset) | Song.rep.final | Non-Cooperative | Cooperative | 0.994 | 0.198 |
| Cooperative breeding classifications from Downing et al (2015) (transition rates based on current study dataset) | Song.rep.final | Non-Cooperative | Cooperative | 0.988 | 0.156 |
| Cooperative breeding classifications from Jetz and Rubenstein (2011) (transition rates based on source dataset) | Song.rep.final | Non-Cooperative | Cooperative | 1 | 0.001 |
| Cooperative breeding classifications from Jetz and Rubenstein (2011) (transition rates based on current study dataset) | Song.rep.final | Non-Cooperative | Cooperative | 0.998 | 0.003 |
| Cooperative breeding classifications from Cockburn (2016) (transition rates based on source dataset) | Song.rep.final | Non-Cooperative | Cooperative | 0.996 | 0.003 |
| Cooperative breeding classifications from Cockburn (2016) (transition rates based on current study dataset) | Song.rep.final | Non-Cooperative | Cooperative | 0.994 | 0.003 |
| Cooperative breeding classifications from Griesser et al (2017) (transition rates based on source dataset) | Song.rep.final | Non-Cooperative | Cooperative | 1 | 0 |
| Cooperative breeding classifications from Griesser et al (2017) (transition rates based on current study dataset) | Song.rep.final | Non-Cooperative | Cooperative | 1 | 0 |
| Cooperative breeding classifications from Dale (2015) (transition rates based on source dataset) | Song.rep.final | Non-Cooperative | Cooperative | 0.952 | 0.137 |
| Cooperative breeding classifications from Dale (2015) (transition rates based on current study dataset) | Song.rep.final | Non-Cooperative | Cooperative | 0.91 | 0.178 |
| Cooperative breeding classifications from Cornwallis (2017) (transition rates based on source dataset) | Song.rep.final | Non-Cooperative | Cooperative | 0.996 | 0.021 |
| Cooperative breeding classifications from Cornwallis (2017) (transition rates based on current study dataset) | Song.rep.final | Non-Cooperative | Cooperative | 0.988 | 0.046 |

**Supplemental Table 5:** Results assessing whether the rate of song repertoire size evolution differs based on the predicted ancestral state of cooperative breeding status across subsets of species, iteratively removing each family with more than 2 species in the analysis. Each subset was run for 100 simulations using the Brownie algorithm (brownie.lite, R package: phytools). A likelihood-ratio test was used to calculate *p*-values.

| Song Trait | Higher rate of evolution in state: | Lower rate of evolution in state: | Fraction of simulations following trend | Brownie *p*-value | Number of Species In Subset Without Removed Family | Removed Family |
|---|---|---|---|---|---|---|
| Song.rep.final | Non-Cooperative | Cooperative | 1 | 0.03 | 209 | None |
| Song.rep.final | Non-Cooperative | Cooperative | 1 | 0.025 | 202 | Cardinalidae |
| Song.rep.final | Non-Cooperative | Cooperative | 1 | 0.025 | 207 | Certhiidae |
| Song.rep.final | Non-Cooperative | Cooperative | 0.99 | 0.044 | 207 | Corvidae |
| Song.rep.final | Non-Cooperative | Cooperative | 1 | 0.033 | 162 | Emberizidae |
| Song.rep.final | Non-Cooperative | Cooperative | 1 | 0.03 | 206 | Estrildidae |
| Song.rep.final | Non-Cooperative | Cooperative | 1 | 0.031 | 203 | Fringillidae |
| Song.rep.final | Non-Cooperative | Cooperative | 1 | 0.018 | 206 | Hirundinidae |
| Song.rep.final | Non-Cooperative | Cooperative | 1 | 0.027 | 193 | Icteridae |
| Song.rep.final | Non-Cooperative | Cooperative | 0.98 | 0.035 | 205 | Mimidae |
| Song.rep.final | Non-Cooperative | Cooperative | 1 | 0.025 | 206 | Motacillidae |
| Song.rep.final | Non-Cooperative | Cooperative | 1 | 0.031 | 203 | Muscicapidae |
| Song.rep.final | Non-Cooperative | Cooperative | 0.99 | 0.033 | 197 | Paridae |
| Song.rep.final | Non-Cooperative | Cooperative | 0.99 | 0.02 | 182 | Parulidae |
| Song.rep.final | Non-Cooperative | Cooperative | 0.99 | 0.03 | 207 | Reguliidae |
| Song.rep.final | Non-Cooperative | Cooperative | 1 | 0.058 | 193 | Sylviidae |
| Song.rep.final | Non-Cooperative | Cooperative | 0.98 | 0.032 | 196 | Troglodytidae |
| Song.rep.final | Non-Cooperative | Cooperative | 1 | 0.033 | 195 | Turdidae |
| Song.rep.final | Non-Cooperative | Cooperative | 0.99 | 0.038 | 207 | Viduidae |
| Song.rep.final | Non-Cooperative | Cooperative | 1 | 0.028 | 198 | Vireonidae |

**Supplemental Table 6:** Results assessing whether the rate of song repertoire size evolution differs based on the predicted ancestral state of several multi-state traits: (A) group size, with four possible values, “asocial”, “pair”, “small groups”, and “large groups” (Griesser et al 2023), (B) territoriality, with three levels of classification, across 500 simulations using the Brownie algorithm (brownie.lite, R package: phytools), (C) combined territoriality strength and cooperative breeding across 1500 simulations, (D) social system, with three possible values (with the two non-familial groups merged, Griesser et al 2017), and (E) social system, with four possible values, “Cooperative, Familial”, “Cooperative, Non-familial”, “Non-cooperative, Familial”, and “Non-cooperative, Non-familial” (Griesser et al 2017). The columns “Higher Rate” and “Lower Rate” show which member of each pair of states was found to evolve at a higher and lower rate (respectively) the majority of the time, and the “Fraction of Simulations” columns indicate the frequency out of 500 simulations in which that pattern of higher- and lower-rate states was observed (those comparisons that were consistent in direction over at least 95% of simulations are highlighted in green). Overall, we found evidence that song repertoire size evolves at significantly different rates in lineages with different group sizes (Brownie likelihood-ratio test *p*-value <0.001; n = 127 species), territoriality (Brownie likelihood-ratio test *p*-value = 0.012; n = 216 species), interactions between cooperative breeding and territory strength (Brownie likelihood-ratio test *p*-value = 0.011; n = 152 species), and both methods of categorizing social system (Brownie likelihood-ratio test 3-state *p*-value = 0.001, 4-state *p*-value = 0.002; both n = 156 species). See Extended Data Figure 2.

| | Evolution of song repertoire size | | |
|---|---|---|---|
| Social feature name | Higher Rate | Lower Rate | Fraction of Simulations |
| (A) grouping (Griesser et al 2023) | pair | asocial | 1 |
| | pair | large_groups | 1 |
| | pair | small_groups | 1 |
| | large_groups | asocial | 0.998 |
| | large_groups | small_groups | 0.988 |
| | asocial | small_groups | 0.824 |
| (B) Territory (Tobias et al, 2016) | Seasonal or weak territoriality (2) | Year-round territoriality (3) | 1 |
| | No territoriality (1) | Year-round territoriality (3) | 1 |
| | No territoriality (1) | Seasonal or weak territoriality (2) | 0.996 |
| (C) Cooperative Breeding x Weak vs Strong Territoriality (this paper) | Noncooperative_Weak | Cooperative_Strong | 0.996 |
| | Noncooperative_Strong | Cooperative_Strong | 0.995 |
| | Noncooperative_Weak | Noncooperative_Strong | 0.987 |
| | Cooperative_Weak | Cooperative_Strong | 0.928 |
| | Noncooperative_Weak | Cooperative_Weak | 0.679 |
| | Noncooperative_Strong | Cooperative_Weak | 0.586 |
| (D) “social system” three-state category from Griesser et al (2017) | Non-familial | Cooperative, Familial | 0.998 |
| | Non-familial | Non-cooperative, Familial | 0.978 |
| | Non-cooperative, Familial | Cooperative, Familial | 0.64 |
| (E) “social system including nonkin-coop” four-state category from Griesser et al (2017) | Non-cooperative, Non-familial | Cooperative, Familial | 0.998 |
| | Non-cooperative, Non-familial | Non-cooperative, Familial | 0.988 |
| | Non-cooperative, Non-familial | Cooperative, Non-familial | 0.89 |
| | Non-cooperative, Familial | Cooperative, Familial | 0.694 |
| | Cooperative, Non-familial | Cooperative, Familial | 0.658 |
| | Non-cooperative, Familial | Cooperative, Non-familial | 0.558 |

**Supplemental Table 7:** Overlap of stochastic character map simulations of cooperative breeding and female song (data in HighConfidence_Coop and FemaleSong_Agg01 columns in Supplemental Datasets 2 and 3) using a jackknife analysis in which each family was removed from the datasets in turn. Each subset was run for 50 real simulations, 200 randomized-data simulations.

| Removed family | Number of species in subset without removed family | Empirical *p*-value | Degree of association between traits | | | |
|---|---|---|---|---|---|---|
| | | | Non-cooperative breeding co-occurrence with female song absence | Non-cooperative breeding co-occurrence with female song presence | Cooperative breeding co-occurrence with female song absence | Cooperative breeding co-occurrence with female song presence |
| None | 1041 | <0.005 | 0.28 | 0.26 | 0.47 | 1 |
| Acanthizidae | 1020 | 0.005 | 0.325 | 0.255 | 0.37 | 1 |
| Alaudidae | 992 | 0.025 | 0.315 | 0.175 | 0.59 | 1 |
| Campephagidae | 1025 | <0.005 | 0.32 | 0.23 | 0.295 | 1 |
| Cardinalidae | 1028 | <0.005 | 0.235 | 0.185 | 0.51 | 1 |
| Certhiidae | 1038 | 0.015 | 0.22 | 0.315 | 0.51 | 1 |
| Cinclosomatidae | 1038 | <0.005 | 0.255 | 0.3 | 0.565 | 1 |
| Cisticolidae | 974 | <0.005 | 0.54 | 0.07 | 0.6 | 1 |
| Climacteridae | 1036 | 0.01 | 0.22 | 0.265 | 0.56 | 1 |
| Colluricinclidae | 1036 | 0.005 | 0.28 | 0.3 | 0.49 | 1 |
| Corvidae | 1027 | 0.015 | 0.29 | 0.325 | 0.54 | 1 |
| Cracticidae | 1033 | 0.005 | 0.245 | 0.26 | 0.635 | 1 |
| Dasyornithidae | 1038 | 0.02 | 0.225 | 0.275 | 0.415 | 1 |
| Dicruridae | 1029 | <0.005 | 0.215 | 0.175 | 0.44 | 1 |
| Emberizidae | 979 | 0.02 | 0.275 | 0.22 | 0.585 | 1 |
| Estrildidae | 1017 | <0.005 | 0.195 | 0.315 | 0.505 | 1 |
| Eupetidae | 1036 | 0.005 | 0.265 | 0.205 | 0.435 | 1 |
| Fringillidae | 1004 | <0.005 | 0.315 | 0.205 | 0.22 | 1 |
| Hirundinidae | 1036 | 0.01 | 0.225 | 0.275 | 0.495 | 1 |
| Icteridae | 979 | 0.035 | 0.25 | 0.25 | 0.495 | 1 |
| Laniidae | 1031 | 0.01 | 0.24 | 0.22 | 0.455 | 1 |
| Malaconotidae | 1003 | 0.005 | 0.215 | 0.165 | 0.545 | 1 |
| Maluridae | 1027 | 0.01 | 0.225 | 0.315 | 0.385 | 0.995 |
| Meliphagidae | 1003 | <0.005 | 0.23 | 0.235 | 0.49 | 1 |
| Mimidae | 1037 | 0.01 | 0.285 | 0.19 | 0.535 | 1 |
| Monarchidae | 1026 | 0.005 | 0.2 | 0.29 | 0.325 | 1 |
| Motacillidae | 1033 | 0.015 | 0.21 | 0.21 | 0.44 | 1 |
| Muscicapidae | 974 | <0.005 | 0.21 | 0.305 | 0.52 | 1 |
| Nectariniidae | 1024 | <0.005 | 0.12 | 0.275 | 0.445 | 1 |
| Oriolidae | 1033 | 0.005 | 0.29 | 0.175 | 0.475 | 1 |
| Orthonychidae | 1038 | 0.02 | 0.24 | 0.29 | 0.49 | 1 |
| Pachycephalidae | 1036 | <0.005 | 0.25 | 0.3 | 0.395 | 1 |
| Paradisaeidae | 1016 | 0.005 | 0.335 | 0.17 | 0.605 | 1 |
| Pardalotidae | 1038 | <0.005 | 0.28 | 0.205 | 0.42 | 1 |
| Paridae | 1034 | 0.015 | 0.225 | 0.275 | 0.54 | 1 |
| Parulidae | 1023 | 0.005 | 0.19 | 0.31 | 0.44 | 1 |
| Passeridae | 1031 | <0.005 | 0.325 | 0.16 | 0.45 | 1 |
| Petroicidae | 1031 | <0.005 | 0.345 | 0.2 | 0.48 | 1 |
| Platysteiridae | 1035 | 0.02 | 0.25 | 0.255 | 0.535 | 1 |
| Ploceidae | 1019 | <0.005 | 0.285 | 0.24 | 0.315 | 1 |
| Prunellidae | 1038 | 0.005 | 0.19 | 0.275 | 0.435 | 1 |
| Ptilonorhynchidae | 1031 | 0.005 | 0.175 | 0.275 | 0.615 | 1 |
| Pycnonotidae | 1033 | <0.005 | 0.255 | 0.19 | 0.415 | 1 |
| Reguliidae | 1038 | 0.01 | 0.17 | 0.25 | 0.455 | 1 |
| Sittidae | 1033 | <0.005 | 0.37 | 0.115 | 0.4 | 1 |
| Sturnidae | 1021 | <0.005 | 0.315 | 0.21 | 0.285 | 1 |
| Sylviidae | 1009 | <0.005 | 0.235 | 0.23 | 0.495 | 1 |
| Thraupidae | 1011 | 0.005 | 0.245 | 0.215 | 0.585 | 1 |
| Timaliidae | 993 | 0.01 | 0.1 | 0.35 | 0.74 | 1 |

**Supplemental Table 8:** Results of overlapping simulations of ancestral character history for cooperative breeding and female song using alternative phylogenies, alternative methods of determining cooperative breeding, and cooperative breeding classifications based on individual sources, with transition rates determined based on either our full set of species or just the species present in that source, as noted. For each analysis, we ran 500 ancestral character mapping simulations with the real data and 500 with shuffled (randomized) data for comparison, except where otherwise noted. Empirical *p*-value indicates the *p*-value from our adapted Huelsenbeck et al (2003) method. Cells highlighted in yellow indicate significance at $p < 0.05$. Across the Trait A-Trait B stochastic character map pairs based on the real data, we found the median proportion of the phylogeny that each combination of traits occupied, then approximated the degree of association between trait states by calculating the fraction of randomized stochastic character map pairs that had a proportion of the tree occupying a given trait combination below the median of the real simulations. Cells highlighted in blue indicate that that trait combination was greater than the median of real simulations in <5% of randomized simulations (and thus the trait combination is less common than expected if traits were independent). Cells highlighted in red indicate that >95% of randomized simulations were less than the median of the real simulations (thus the trait combination is more common than expected if traits were independent). Colors in the first column follow Supplemental Table 4. For detailed breakdowns of the number of species in each state for each cooperative breeding classification method, see Supplemental Table 9.

| Non-cooperative (0) vs. cooperative (1) data or phylogeny type | Number of species | Empirical *p*-value | Degree of association between Trait A (female song absent versus female song present) and Trait B (non-cooperative versus cooperative breeding) | | | |
|---|---|---|---|---|---|---|
| | | | Female song absent co-occurrence with non-cooperative breeding | Female song present co-occurrence with non-cooperative breeding | Female song absent co-occurrence with cooperative breeding | Female song present co-occurrence with cooperative breeding |
| Alternate consensus tree (mean.edge, if.absent = "ignore") | 1041 | 0.008 | 0.228 | 0.224 | 0.482 | 1 |
| Multiple independent trees (200 trees, 20 simulations per tree) | 1041 | 0.00075 | 0.462 | 0.131 | 0.514 | 1 |
| Alternate cooperative breeding classification method: MeanCoopTie2Coop | 1075 | 0.018 | 0.298 | 0.182 | 0.568 | 1 |
| Alternate cooperative breeding classification method: MeanCoopOmitTies | 1066 | 0.01 | 0.268 | 0.168 | 0.616 | 1 |
| Alternate cooperative breeding classification method: AnyCoopEqualsCoop | 1075 | 0.016 | 0.346 | 0.192 | 0.376 | 1 |
| Alternate cooperative breeding classification method: AnyNoncoopEqualsNoncoop | 1075 | 0.01 | 0.304 | 0.182 | 0.376 | 1 |
| Alternate cooperative breeding classification method: MeanCoopTie2Noncoop | 1075 | 0.012 | 0.254 | 0.136 | 0.734 | 1 |
| Classifications from Cockburn 2006 (transition rates based on source dataset) | 751 | 0.002 | 0.576 | 0.126 | 0.328 | 0.996 |
| Classifications from Cockburn 2006 (transition rates based on current study dataset) | 751 | 0.004 | 0.546 | 0.128 | 0.314 | 1 |
| Classifications from Griesser et al 2017 (transition rates based on source dataset) | 468 | 0.008 | 0.75 | 0.012 | 0.66 | 1 |

| | | | | | | |
|---|---|---|---|---|---|---|
| Classifications from Griesser et al 2017 (transition rates based on current study dataset) | 468 | 0.004 | 0.602 | 0.004 | 0.912 | 1 |
| Classifications from Dale 2015 (transition rates based on source dataset) | 785 | 0.01 | 0.31 | 0.154 | 0.482 | 0.996 |
| Classifications from Dale 2015 (transition rates based on current study dataset) | 785 | 0.004 | 0.304 | 0.184 | 0.532 | 0.998 |
| Classifications from Cornwallis 2017 (transition rates based on source dataset) | 731 | 0.008 | 0.312 | 0.448 | 0.438 | 1 |
| Classifications from Cornwallis 2017 (transition rates based on current study dataset) | 731 | 0.01 | 0.316 | 0.486 | 0.314 | 0.998 |
| Classifications from Jetz and Rubenstein 2011 (transition rates based on source dataset) | 1058 | 0.04 | 0.222 | 0.092 | 0.848 | 1 |
| Classifications from Jetz and Rubenstein 2011 (transition rates based on current study dataset) | 1058 | 0.038 | 0.208 | 0.144 | 0.764 | 1 |
| Classifications from Biagolini et al 2017 (transition rates based on source dataset) | 114 | 0.16 | 0.512 | 0.228 | 0.53 | 0.974 |
| Classifications from Biagolini et al 2017 (transition rates based on current study dataset) | 114 | 0.238 | 0.48 | 0.196 | 0.648 | 0.936 |
| Classifications from Downing et al 2015 (transition rates based on source dataset) | 105 | 0.262 | 0.574 | 0.13 | 0.822 | 0.97 |
| Classifications from Downing et al 2015 (transition rates based on current study dataset) | 105 | 0.268 | 0.542 | 0.136 | 0.856 | 0.954 |

**Supplemental Table 9:** Species counts for each combination of the social system classification from the current study and individual sources used to generate the database, separated by female song presence or absence based on the classification in the current study. Color scheme is the same as Supplemental Table 4.

| Column name | Social System State | Female Song Absent | Female Song Present |
|---|---|---|---|
| HighConfidence_Coop | Non-cooperative | 358 | 558 |
| | Cooperative | 24 | 101 |
| MeanCoopTie2Noncoop | Non-cooperative | 366 | 572 |
| | Cooperative | 29 | 108 |
| MeanCoopTie2Coop | Non-cooperative | 361 | 568 |
| | Cooperative | 34 | 112 |
| MeanCoopOmitTies | Non-cooperative | 361 | 568 |
| | Cooperative | 29 | 108 |
| AnyCoopEqualsCoop | Non-cooperative | 353 | 538 |
| | Cooperative | 42 | 142 |
| AnyNoncoopEqualsNoncoop | Non-cooperative | 374 | 589 |
| | Cooperative | 21 | 91 |
| Cooperative breeding classifications from Biagolini et al (2017) | Non-cooperative | 26 | 71 |
| | Cooperative | 2 | 15 |
| Cooperative breeding classifications from Downing et al (2015) | Non-cooperative | 26 | 62 |
| | Cooperative | 2 | 15 |
| Cooperative breeding classifications from Jetz and Rubenstein (2011) | Non-cooperative | 101 | 191 |
| | Cooperative | 6 | 16 |
| Cooperative breeding classifications from Cockburn (2016) | Non-cooperative | 258 | 383 |
| | Cooperative | 24 | 86 |
| Cooperative breeding classifications from Griesser et al (2017) | Non-cooperative | 140 | 231 |
| | Cooperative | 22 | 75 |
| Cooperative breeding classifications from Dale (2015) | Non-cooperative | 268 | 382 |
| | Cooperative | 37 | 98 |
| Cooperative breeding classifications from Cornwallis (2017) | Non-cooperative | 279 | 411 |
| | Cooperative | 5 | 36 |
| social_system_incl_nk_coop_Griesser2017 | Cooperative/Familial | 16 | 71 |
| | Noncooperative/Familial | 43 | 76 |
| | Cooperative/Nonfamilial | 6 | 4 |
| | Noncooperative/Nonfamilial | 97 | 155 |

**Supplemental Table 10**: Results of overlapping simulated ancestral character histories for binary social variables with those of cooperative breeding. For each pair of traits, we ran 500 ancestral character mapping simulations with the real data and 500 with shuffled data (i.e. with tip states in a randomized order) for comparison. We used methods adapted from [23] to estimate an empirical *p*-value (yellow indicates significance at $p < 0.05$). Degree of association between states: Across the pairs of stochastic character maps generated from the real social trait and cooperative breeding data, we found the median proportion of the phylogeny that each combination of traits occupied, then found the fraction of randomized stochastic character map pairs that had a proportion of the tree occupying a given trait combination below the median of the real simulations. Cells highlighted in blue indicate that, in <5% of the randomized-data simulations, the fraction of the tree occupying the indicated state combination was less than the median of simulations based on real data (and thus the state combination is less common than expected if traits were evolving independently). Cells highlighted in red indicate that, in >95% of randomized-data simulations, the state combination occupied a fraction of the tree less than the median of the real simulations (thus the trait combination is more common than expected if traits were evolving independently).

| | | | Degree of association between states of Trait A (non-cooperative breeding versus cooperative breeding) and Trait B | | | |
|---|---|---|---|---|---|---|
| Social Trait (Trait B) | Number of species | Empirical p-value | Non-cooperative breeding co-occurrence with Social Trait = 0 | Cooperative breeding co-occurrence with Social Trait = 0 | Non-cooperative breeding co-occurrence with Social Trait = 1 | Cooperative breeding co-occurrence with Social Trait = 1 |
| Asocial (0) vs. social groups (1) | 441 | 0.006 | 0.78 | 0.104 | 0.01 | 1 |
| Asocial or pair (0) vs larger groups (1) | 441 | 0.006 | 0.914 | 0.088 | 0.004 | 1 |
| Asocial, pair, or small group (0) vs. large group (1) | 441 | <0.002 | 0.016 | 1 | 0.664 | 0.314 |
| Social bonds lasting one season or less (0) vs longer than one breeding season (1) | 472 | <0.002 | 0.018 | 0.002 | 0.56 | 1 |
| Non-familial (0) vs. familial living (1) | 1159 | <0.002 | 0.372 | 0.024 | 0 | 1 |
| Non-colonial (0) vs. colonial living (1) | 472 | 0.036 | 0.042 | 1 | 0.624 | 0.584 |
| Weak or no territoriality (0) vs. strong territoriality (1) | 2012 | <0.002 | 0.41 | 0.686 | 0 | 1 |
| Weak, seasonal, or no territoriality (0) vs. year-round territoriality (1) | 4143 | <0.002 | 0 | 1 | 1 | 1 |
| Monogamy (0) vs. polygyny (1) | 709 | 0.024 | 0.03 | 1 | 0.62 | 0.49 |
| One or two caretakers (0) vs. more than two caretakers (1) | 472 | <0.002 | 0.094 | 0 | 0 | 1 |

**Supplemental Table 11:** Overlap of stochastic character map simulations of female song and four-state social traits. The degree of association between trait states was obtained by calculating the fraction of randomized stochastic character map pairs that had a proportion of the tree occupying a given trait combination below the median value from the real simulations. A) Combined classification of breeding system including kin living and cooperative breeding ("social system including nonkin-coop" from Griesser et al 2017). The empirical *p*-value comparing the overlap between estimated ancestral states of these traits over 1500 simulations is 0.0173, indicating that breeding system and female song co-occur significantly more than expected by chance. Cells highlighted in blue indicate that that trait combination was less than the median of real simulations in <5% of randomized simulations (and thus the trait combination is less common than expected if traits were independent). Cells highlighted in red indicate that >95% of randomized simulations were less than the median of the real simulations (thus the trait combination is more common than expected if traits were independent). Data correspond to Figure 3C. B) Group size classification ("grouping" from Griesser et al 2023). The empirical *p*-value over 1500 simulations is 0.267, indicating that female song is not significantly associated with group size. Sample sizes: (A) n = 468 species, (B) n = 255 species. Data correspond to Extended Data Figure 3B.

| | Degree of association between traits | Female song state | |
|---|---|---|---|
| | Social system state | Absent | Present |
| A | Familial/Cooperative | 0.607 | 1 |
| A | Familial/Non-cooperative | 0.551 | 0.196 |
| A | Non-familial/Cooperative | 0.342 | 0.028 |
| A | Non-familial/Non-cooperative | 0.825 | 0.024 |
| B | Asocial | 0.622 | 0.464 |
| B | Pair | 0.505 | 0.172 |
| B | Small groups | 0.37 | 0.891 |
| B | Large groups | 0.719 | 0.355 |

**Supplemental Table 12**: Distribution of species across cooperative breeding and female song states within different territorial contexts. The table shows raw species counts in our dataset for each combination of traits using two territorial classification schemes: (A) weak versus strong territoriality based on defensive behavior intensity, and (B) whether or not the species was territorial year-round. Cell shading indicates deviation from the expected counts under independence, calculated as (observed-expected)/expected, where blue indicates fewer species than expected and red indicates more species than expected if female song and cooperative breeding were distributed independently within each territorial category. These raw counts do not account for shared ancestral history, but the trends are consistent with the results of our phylogenetic comparative analyses. In each territory category, female song presence and cooperative breeding co-occurred more often than expected (ranging from 98% more than expected in the weak or no territoriality category to 0.9% more than expected in the year-round territoriality category). In both territory classification schemes, female song presence and cooperative breeding were more common in the more territorial category and co-occurred slightly more often than expected by chance, whereas in the less territorial category female song presence and cooperative breeding were less common overall but co-occurred much more often than expected by chance.

| | | | Non-cooperative | Cooperative |
|---|---|---|---|---|
| A | Weak or no territoriality | Female Song Absent | 101 | 3 |
| | | Female Song Present | 56 | 10 |
| | Strong territoriality | Female Song Absent | 172 | 20 |
| | | Female Song Present | 425 | 88 |
| B | Weak, seasonal, or no territoriality | Female Song Absent | 322 | 15 |
| | | Female Song Present | 277 | 23 |
| | Year-round territoriality | Female Song Absent | 36 | 9 |
| | | Female Song Present | 281 | 78 |

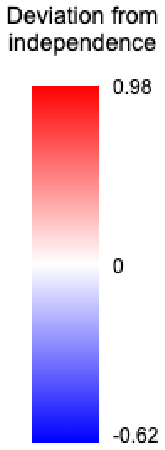

**Supplemental Table 13:** Tests for sampling biases in female song and cooperative breeding data availability. We tested whether data availability for female song presence/absence and cooperative breeding classification was biased with respect to various species traits that could confound evolutionary analyses. For each bias type, we tested the null hypothesis that data availability was independent of the trait in question. Chi-squared tests were used for categorical traits and Welch's t-tests for continuous traits. Significant biases ($p < 0.05$) indicate that species with certain trait values are over- or under-represented in our dataset. Geographic bias was tested using biogeographic regions from Jetz & Rubenstein (2011), with Holarctic (Nearctic + Palearctic) compared to tropical regions. Sexual dichromatism values were obtained from Dale et al. (2015) and sexual dimorphism from wing measurements in AVONET (Tobias et al. 2022). These detected biases were addressed through downsampling procedures (Table S14) to ensure the robustness of our main findings.

| Bias Type | Hypothesis | Test | Test statistic | df | p-value | Result |
|---|---|---|---|---|---|---|
| Cooperative breeding | Cooperative breeding species more likely to have female song data | Chi-squared | $\chi^2 = 2.18$ | 1 | 0.14 | No bias in FS data |
| Coloniality | Colonial species more likely to be studied | Chi-squared<br>Chi-squared | $\chi^2 = 0.01$<br>$\chi^2 = 2.06$ | 1<br>1 | 0.91<br>0.15 | No bias in FS data<br>No bias in CB data |
| Polygyny | Polygynous species more likely to be studied | Chi-squared<br>Chi-squared | $\chi^2 = 2.143$<br>$\chi^2 = 0.46$ | 1<br>1 | 0.143<br>0.498 | No bias in FS data<br>No bias in CB data |
| Geographic | Holarctic species more likely to be studied | Chi-squared<br>Chi-squared | $\chi^2 = 56.80$<br>$\chi^2 = 0.74$ | 1<br>1 | <0.001<br>0.785 | **Bias detected in FS data**<br>No bias in CB data |
| Geographic x Cooperative breeding | Female song may be more or less likely to be studied based on the interaction between their geographic region and cooperative breeding status | Chi-squared | $\chi^2 = 56.92$ | 3 | <0.001 | **Bias detected in FS data** (Post-hoc tests found that non-cooperative holarctic species and cooperative tropical species are both overrepresented in the female song data) |
| Territoriality (Weak/Strong) | More territorial more likely to be studied | Chi-squared<br>Chi-squared | $\chi^2 = 58.67$<br>$\chi^2 = 31.49$ | 1<br>1 | <0.001<br><0.001 | **Bias detected in FS data**<br>**Bias detected in CB data** |
| Territory (Year-round) | Year-round territorial more likely to be studied | Chi-squared<br>Chi-squared | $\chi^2 = 89.22$<br>$\chi^2 = 39.18$ | 1<br>1 | <0.001<br><0.001 | **Bias detected in FS data**<br>**Bias detected in CB data** |
| Sexual dichromatism | Dichromatic species more likely to have FS data | t-test | $t = 3.46$ | 1789 | <0.001 | **Bias detected in FS data** |
| Sexual dimorphism | Dimorphic species more likely to have FS data | t-test | $t = -2.77$ | 1759 | 0.006 | **Bias detected in FS data** |

**Supplemental Table 14:** Stratified downsampling calculations to correct for sampling biases in the comparative dataset. For each identified bias (Table S13), we calculated the number of species to remove to achieve balanced representation across groups. Current proportions show the fraction of species with complete female song and cooperative breeding data within each group. Target proportions represent the lower of the two groups being compared, ensuring conservative bias correction. For geographic biases, we performed two complementary corrections: removing overrepresented Holarctic non-cooperative species, and reducing the excess of tropical cooperative species among those with data. Although female song data availability was not significantly greater for cooperative breeding species globally, we performed an additional adjustment of the global cooperative breeding rate with female song data to match the population rate. All downsampling was performed independently 500 times.

| Bias Category | Group to Downsample | Species to Remove | Current proportion of [Group to Downsample] with FS & CB data available | Target Proportion | Justification |
|---|---|---|---|---|---|
| Geographic | Holarctic non-cooperative | 83 | 35.3% (209/592) | 21.3% | Match tropical non-cooperative proportion |
| Geographic | Tropical cooperative | 24 | 13.9% (112/804) | 11.3% | Match overall tropical cooperative breeding rate |
| Cooperative breeding | Global cooperative | 15 | 12.0% (125/1041) | 10.7% | Match population cooperative breeding rate |
| Territoriality (Weak/Strong) | Strong territoriality | 266 | 45.2% (705/1560) | 28.1% | Match weak territoriality proportion |
| Territoriality (Year-round) | Year-round territoriality | 155 | 32.4% (404/1246) | 20.0% | Match seasonal/weak/no territory proportion |
| Sexual dichromatism | Continuous metric: preferentially downsample more dichromatic species | 167 | Remove species to approximately reduce the mean log(abs(plumage dimorphism)) (pre-normalization) of the subset for which we do have female song data (1.02) to the mean of the group for which we do not have data (0.86) | | |
| Sexual dimorphism | Continuous metric: preferentially downsample more dimorphic species | 71 | Remove species to approximately reduce the mean % abs(wing dimorphism) (pre-normalization) of the subset for which we do have female song data (1.11) to the mean of the group for which we do not have data (1.04) | | |

**Supplemental Table 15:** Results of phylogenetic generalized linear models using alternative cooperative breeding classifications with female song as the response variable. The alternative cooperative breeding variables were inserted into the best-fit model from the tests performed using our primary cooperative breeding variable, Female Song ~ Cooperative Breeding * Territoriality (Weak vs Strong) + log(Mass) (centered and normalized). Cells display the mean Odds Ratio across 500 bootstrap iterations. Cells are filled if the empirical *p*-value is < 0.05, indicating that the bootstrapped odds ratio confidence interval does not include 1.0, with red cells indicating an odds ratio >1 and blue cells indicating an odds ratio <1. Significance is also indicated by symbols: *** p < 0.001, ** p < 0.01, * p < 0.05, ^ p < 0.1.

| Cooperative Breeding Classifier | Response Variable | Species n | Intercept | Cooperative Breeding | Territoriality | Body Mass | CBxTerr |
|---|---|---|---|---|---|---|---|
| MeanCoopTie2Noncoop | Female Song | 902 | 0.653^ | 4.370* | 3.683*** | 1.411*** | 0.289* |
| MeanCoopTie2Coop | Female Song | 902 | 0.654^ | 5.371*** | 3.567*** | 1.474* | 0.222** |
| MeanCoopOmitTies | Female Song | 895 | 0.631* | 4.655* | 3.642*** | 1.466*** | 0.272* |
| AnyCoopEqualsCoop | Female Song | 902 | 0.534* | 9.486*** | 4.477*** | 1.454*** | 0.110*** |
| AnyNoncoop EqualsNoncoop | Female Song | 902 | 0.663* | 3.612* | 3.637*** | 1.387* | 0.379^ |
| CockburnCoop | Female Song | 650 | 0.597* | 4.334** | 4.952*** | 1.220^ | 0.290* |
| CockburnInferred | Female Song | 864 | 0.648^ | 4.251*** | 3.868*** | 1.491** | 0.256** |
| HighConf_Coop DefaultToCockburnInferred | Female Song | 904 | 0.630* | 4.941*** | 3.787*** | 1.490*** | 0.219** |
| Griesser2017Coop | Female Song | 424 | 0.744 | 2.534 | 3.812*** | 1.776** | 0.541 |
| DaleCoop | Female Song | 681 | 0.664^ | 6.738*** | 4.095*** | 1.198 | 0.136*** |
| CornwallisCoop | Female Song | 634 | 0.707 | 2.112^ | 3.978*** | 1.190 | 1.356 |
| JetzCoopInclCockburn | Female Song | 888 | 0.640* | 4.706*** | 3.906*** | 1.477*** | 0.225* |

**Supplemental Table 16:** Results of phylogenetic generalized linear models using alternative cooperative breeding classifications with cooperative breeding as the response variable. The alternative cooperative breeding variables were inserted into the best-fit model from the tests performed using our primary cooperative breeding variable, Cooperative Breeding ~ Female Song * Territoriality (Weak vs Strong). Cells display the mean Odds Ratio across 500 bootstrap iterations. Cells are filled if the empirical *p*-value is < 0.05, indicating that the bootstrapped odds ratio confidence interval does not include 1.0, with red cells indicating an odds ratio >1 and blue cells indicating an odds ratio <1. Significance is also indicated by symbols: *** p < 0.001, ** p < 0.01, * p < 0.05, ^ p < 0.1.

| Response Variable | Species n | Intercept | Female Song | Territoriality | FSxTerr |
|---|---|---|---|---|---|
| MeanCoopTie2Noncoop | 902 | 0.100** | 5.514** | 5.549** | 0.181** |
| MeanCoopTie2Coop | 902 | 0.099*** | 6.639** | 7.077*** | 0.146** |
| MeanCoopOmitTies | 895 | 0.104*** | 5.621*** | 5.844*** | 0.175*** |
| AnyCoopEqualsCoop | 902 | 0.134*** | 11.343*** | 9.033*** | 0.090*** |
| AnyNoncoopEqualsNoncoop | 902 | 0.115*** | 4.614*** | 4.454*** | 0.225*** |
| CockburnCoop | 650 | 0.099*** | 4.706*** | 4.230*** | 0.206*** |
| CockburnInferred | 864 | 0.179*** | 5.396*** | 5.656*** | 0.174*** |
| HighConf_Coop DefaultToCockburnInferred | 904 | 0.133*** | 6.244*** | 6.103*** | 0.152*** |
| Griesser2017Coop | 424 | 0.196** | 2.707* | 2.564* | 0.339* |
| DaleCoop | 681 | 0.113*** | 7.811*** | 5.390*** | 0.115*** |
| CornwallisCoop | 634 | 0.222^ | 1.493 | 0.994 | 0.757 |
| JetzCoopInclCockburn | 888 | 0.132** | 5.663** | 6.010** | 0.165** |

**Supplemental Table 17:** Best-fit models from a phylogenetic generalized linear modeling analysis that included female song, cooperative breeding, year-round territoriality, and body mass. Significant *p*-values ($p < 0.05$) are shown in bold and indicate that the 95% confidence interval of a predictor's odds ratio does not include 1.0.

| *Predictor (of Female Song)* | *Odds Ratio* | *CI* | *p* | *Predictor (of Cooperative Breeding)* | *Odds Ratio* | *CI* | *p* |
|---|---|---|---|---|---|---|---|
| (Intercept) | 0.77 | 0.51-1.11 | 0.180 | (Intercept) | 0.45 | 0.06-1.42 | 0.332 |
| Cooperative Breeding | 1.41 | 0.91-2.28 | 0.088 | Female Song | 1.44 | 1.15-1.93 | **0.012** |
| Year-round territory | 5.72 | 4.04-8.10 | **<0.002** | Year-round territory | 1.91 | 1.19-3.79 | **<0.002** |
| log(Mass) (normalized | 1.28 | 1.11-1.45 | **0.004** | FemaleSong:Territory | 0.59 | 0.30-1.05 | 0.116 |
| Species | 1041 | | | Species | 1041 | | |
| Model formula: Female Song ~ Cooperative Breeding + Territoriality (Year-round) + log(Mass) | | | | Model formula: Cooperative Breeding ~ Female Song * Territory (Year-round) | | | |

**Supplemental Table 18:** Phylogenetic generalized linear model comparisons of cooperative breeding versus alternative social predictors of female song. We compared cooperative breeding against each social variable using identical model structures with additive predictors (Female Song ~ [Social Variable] + Territoriality + Body Mass). Sample sizes represent species with complete data for both cooperative breeding and the social variable being compared. All ordinal or continuous social variables were standardized (scaled to have a standard deviation of 1 and mean of 0). The odds ratio for each predictor is reported as the mean and 95% CI across 500 bootstrap iterations. The *p*-value is determined empirically based on the fraction of bootstrap iterations where the predictor's odds ratio falls above or below 1.0.

| | | | Results from model using **cooperative breeding** as the predictor (Female Song ~ Cooperative Breeding + Territoriality + Body Mass) | | | | Results from model using the **social variable** as the predictor (Female Song ~ [Social Variable] + Territoriality + Body Mass) | | | |
|---|---|---|---|---|---|---|---|---|---|---|
| Territoriality variable | Social Variable | N Species | CB Model AIC | CB Odds Ratio | CB Odds Ratio 95% CI | CB *p*-value | Social Model AIC | Social Variable Odds Ratio | Social Variable Odds Ratio 95% CI | Social Variable *p*-value |
| Territoriality (Year-Round) | caretakers (normalized) | 258 | 251.3 | 3.15 | 1.11-15.97 | **0.044** | **242.9** | **9.13** | 2.44-33.83 | **0.02** |
| | Familial living | 448 | 461.9 | **1.50** | 0.84-2.97 | 0.2 | 462.0 | 0.78 | 0.49-1.25 | 0.268 |
| | Colonial living | 258 | 251.3 | **3.27** | 1.12-18.82 | **0.028** | 252.5 | 1.22 | 0.62-2.17 | 0.472 |
| | Grouping (normalized) | 243 | 235.3 | **3.31** | 1.19-14.85 | **0.016** | 233.6 | 0.82 | 0.62-1.01 | 0.072 |
| | Social bond duration (normalized) | 258 | 251.3 | **3.38** | 1.20-18.00 | **0.016** | 251.2 | 1.15 | 0.84-1.57 | 0.264 |

**Supplemental Table 19:** Rates of transition between states of binary traits obtained using Ancestral Character Estimation (R Package 'ape', function 'ace') with the Equal Rates (ER) and All Rates Different (ARD) models, calculated using Oscine species only. The *p*-value was obtained using function anova() to compare the ER and ARD models. Bolded values indicate the rates used in simulations. Except where otherwise noted, all rates were calculated using the consensus tree calculated using default settings. See Supplemental Table 1 for "0" and "1" binary states for each trait.

| trait | Rate of transition (Equal Rates; 0→1 = 1→0) | Rate of transition 0→1 (All Rates Different) | Rate of transition 1→0 (All Rates Different) | Log-Likelihood (Equal Rates model) | Log-Likelihood (All Rates Different model) | Likelihood ratio test *p*-value |
|---|---|---|---|---|---|---|
| Griesser2023.Colonial01 | 0.01269992 | **0.01200588** | **0.08386504** | -213.22631 | -186.39854 | <0.001 |
| Griesser2023.MoreThanTwoCaretakers | **0.00963502** | 0.00966078 | 0.0162108 | -164.98296 | -164.52879 | 0.341 |
| Griesser2023.LongSocialBonds | 0.01945464 | **0.04150603** | **0.00849845** | -241.33754 | -230.45623 | <0.001 |
| Griesser2023.GroupsLargerThanPair | 0.01094027 | **0.04238356** | **0.00869378** | -168.51113 | -155.24743 | <0.001 |
| Griesser2023.Asocial0VsSocial1 | 0.00778502 | **0.06909473** | **0.00454807** | -144.41485 | -112.31853 | <0.001 |
| Griesser2023.LargestGroupSizes | 0.03826034 | **0.03400503** | **0.0976376** | -293.0195 | -272.68888 | <0.001 |
| Griesser2017FamilialLiving | 0.05339407 | **0.06052542** | **0.04661725** | -715.94102 | -713.34626 | 0.023 |
| Final.polygyny | 0.01269635 | **0.01199225** | **0.1090912** | -269.17593 | -242.80781 | <0.001 |
| Territory_YearRound | 0.02718114 | **0.02175005** | **0.05177754** | -1806.8906 | -1790.0845 | <0.001 |
| TerritorialityWeakVsStrong | 0.01016844 | **0.02487365** | **0.00718674** | -535.6235 | -520.5321 | <0.001 |
| HighConfidence_Coop | 0.01113287 | **0.01071659** | **0.030594** | -1027.5535 | -1019.6542 | <0.001 |
| FemaleSong_Agg01 | 0.03898356 | **0.06718569** | **0.03828766** | -605.20332 | -597.20892 | <0.001 |
| MeanCoopTie2Noncoop | 0.01274633 | **0.01190063** | **0.05181567** | -1164.7677 | -1137.9685 | <0.001 |
| MeanCoopTie2Coop | 0.01383931 | **0.01343266** | **0.03829614** | -1218.3551 | -1206.3045 | <0.001 |
| MeanCoopOmitTies | 0.01222403 | **0.01174713** | **0.03745765** | -1106.9228 | -1092.8314 | <0.001 |
| AnyCoopEqualsCoop | 0.01745169 | **0.01751687** | **0.03761033** | -1420.5874 | -1412.5426 | <0.001 |
| AnyNoncoopEqualsNoncoop | 0.01143156 | **0.01022155** | **0.06264396** | -1090.1805 | -1048.4043 | <0.001 |
| HighConfidence_Coop (consensus tree with missing edges ignored in mean edge length calculation) | 0.01084228 | **0.01036728** | **0.03159119** | -1027.534 | -1018.734 | <0.001 |
| FemaleSong_Agg01 (consensus tree with missing edges ignored in mean edge length calculation) | 0.03697411 | **0.06447600** | **0.03646347** | -603.8432 | -595.8721 | <0.001 |

**Supplemental Table 20:** Comparing values of standardized (i.e. scaled to have a standard deviation of 1 and center of 0) continuous and ordinal variables across female song and cooperative breeding groups using phylANOVA.

| Variable | Evaluating differences between the states of trait: | Phyl-ANOVA p-value | Mean of group with trait state 0 | SD of group with trait state 0 | N Species with trait state 0 | Mean of group with trait state 1 | SD of group with trait state 1 | N Species with trait state 1 |
|---|---|---|---|---|---|---|---|---|
| abs(Latitude) | Female Song | 0.58038 | 0.273 | 1.120 | 393 | 0.374 | 1.175 | 688 |
| Number of caretakers | Female Song | **0.02552** | -0.326 | 0.340 | 70 | **0.011** | 0.903 | 200 |
| grouping | Female Song | 0.26032 | **0.084** | 1.141 | 64 | -0.168 | 1.056 | 191 |
| Body mass | Female Song | 0.24686 | 0.067 | 1.052 | 397 | 0.251 | 0.965 | 697 |
| Social bond duration | Female Song | **0.0016** | -0.460 | 1.331 | 70 | **0.127** | 0.872 | 200 |
| Plumage dichromatism | Female Song | 0.07304 | 0.272 | 0.999 | 397 | -0.010 | 1.015 | 697 |
| Wing dimorphism | Female Song | **0.04788** | **0.257** | 1.191 | 375 | -0.048 | 0.876 | 660 |
| abs(Latitude) | Cooperative Breeding | 0.86344 | 0.006 | 1.020 | 3847 | -0.030 | 0.891 | 460 |
| Number of caretakers | Cooperative Breeding | **2.00E-05** | -0.256 | 0.198 | 402 | **1.471** | 2.054 | 70 |
| grouping | Cooperative Breeding | 0.90294 | -0.007 | 1.087 | 371 | 0.042 | 0.284 | 70 |
| Body mass | Cooperative Breeding | **0.02648** | -0.040 | 0.982 | 3904 | **0.363** | 1.119 | 469 |
| Social bond duration | Cooperative Breeding | **0.02124** | -0.106 | 1.041 | 402 | **0.615** | 0.365 | 70 |
| Plumage dichromatism | Cooperative Breeding | 0.06518 | 0.050 | 1.010 | 3848 | -0.296 | 0.898 | 464 |
| Wing dimorphism | Cooperative Breeding | 0.36196 | 0.023 | 1.019 | 3547 | -0.173 | 0.880 | 438 |

**Supplemental Table 21:** Best phylogenetic generalized linear model predicting female song after stepwise predictor addition, using territory strength as the territorial categorization. Predictors that were tested but not retained through the stepwise iterative process include latitude (distance from equator), geographic region, migration, and social mating system (polygyny). *p*-values less than 0.05 are in bold.

| *Predictors (of Female Song)* | *Odds Ratio* | *CI* | *p* |
|---|---|---|---|
| (Intercept) | 0.62 | 0.38-1.00 | **0.048** |
| HighConfidence_Coop | 5.87 | 4.01-8.87 | **0.004** |
| TerritorialityWeakVsStrong | 4.73 | 3.06-7.28 | **<0.002** |
| logMass_normalized | 1.31 | 1.05-1.63 | **0.024** |
| PercentAbsLogWingDimorphism_normalized | 0.83 | 0.71-0.97 | **0.008** |
| logMaleFemalePlumageDiffAbs_normalized | 0.85 | 0.71-1.00 | 0.056 |
| HighConfidence_Coop:TerritorialityWeakVsStrong | 0.22 | 0.16-0.36 | **0.012** |
| Species | 826 | | |

**Supplemental Table 22:** Best phylogenetic generalized linear model predicting cooperative breeding after stepwise predictor addition, using territory strength as the territorial categorization. Predictors that were tested but not retained through the stepwise iterative process include latitude (distance from equator), geographic region, wing dimorphism, plumage dimorphism, migration, and social mating system (polygyny). *p*-values less than 0.05 are in bold.

| *Predictors (of Cooperative Breeding)* | *Odds Ratio* | *CI* | *p* |
|---|---|---|---|
| (Intercept) | 0.02 | 0.01-0.10 | **<0.002** |
| FemaleSong_Agg01 | 11.74 | 3.42-27.39 | **<0.002** |
| TerritorialityWeakVsStrong | 2.41 | 0.85-5.83 | 0.080 |
| Griesser2017FamilialLiving | 14.61 | 5.28-35.33 | **<0.002** |
| FemaleSong_Agg01:TerritorialityWeakVsStrong | 0.07 | 0.02-0.29 | **<0.002** |
| Species | 407 | | |

**Supplemental Table 23:** Best phylogenetic generalized linear model predicting female song after stepwise predictor addition, using year-round territoriality as the territorial categorization. Predictors that were tested but not retained through the stepwise iterative process include plumage dimorphism, migration, and social mating system (polygyny). *p*-values less than 0.05 are in bold.

| *Predictors (of Female Song)* | *Odds Ratio* | *CI* | *p* |
|---|---|---|---|
| (Intercept) | 0.73 | 0.45-1.12 | 0.148 |
| HighConfidence_Coop | 1.49 | 1.02-2.16 | **0.040** |
| Territory_YearRound | 7.02 | 4.94-10.45 | **<0.002** |
| logMass_normalized | 1.52 | 1.31-1.79 | **<0.002** |
| abs_Latitude_normalized | 1.23 | 1.05-1.42 | **0.012** |
| PercentAbsLogWingDimorphism_normalized | 0.87 | 0.75-0.98 | **0.020** |
| GeographicRegion_JetzTropical | 0.67 | 0.46-0.99 | **0.044** |
| Species | 961 | | |

**Supplemental Table 24:** Best phylogenetic generalized linear model predicting cooperative breeding after stepwise predictor addition, using year-round territoriality as the territorial categorization. Predictors that were tested but not retained through the stepwise iterative process include latitude (distance from equator), geographic region, wing dimorphism, plumage dimorphism, migration, and social mating system (polygyny). *p*-values less than 0.05 are in bold.

| *Predictors (of Cooperative Breeding)* | *Odds Ratio* | *CI* | *p* |
|---|---|---|---|
| (Intercept) | 0.04 | 0.01-0.14 | **<0.002** |
| FemaleSong_Agg01 | 1.93 | 1.00-4.12 | 0.052 |
| Territory_YearRound | 2.67 | 0.87-7.14 | 0.072 |
| Griesser2017FamilialLiving | 10.62 | 4.48-24.33 | **<0.002** |
| FemaleSong_Agg01:Territory_YearRound | 0.32 | 0.08-1.05 | 0.064 |
| Species | 448 | | |

**Supplemental Table 25:** Species counts with each combination of trait states for traits geographic region (based on Jetz and Rubenstein 2011), cooperative breeding, and female song.

| | | Female Song | |
|---|---|---|---|
| Geographic Region | Cooperative Breeding | Absent | Present |
| Holarctic | non-cooperative | 69 | 140 |
| Tropical | non-cooperative | 280 | 412 |
| Holarctic | cooperative | 3 | 8 |
| Tropical | cooperative | 20 | 92 |

**Supplemental Figure 1:** Directed acyclic graph plots of all model structures tested in all phylopath analyses.

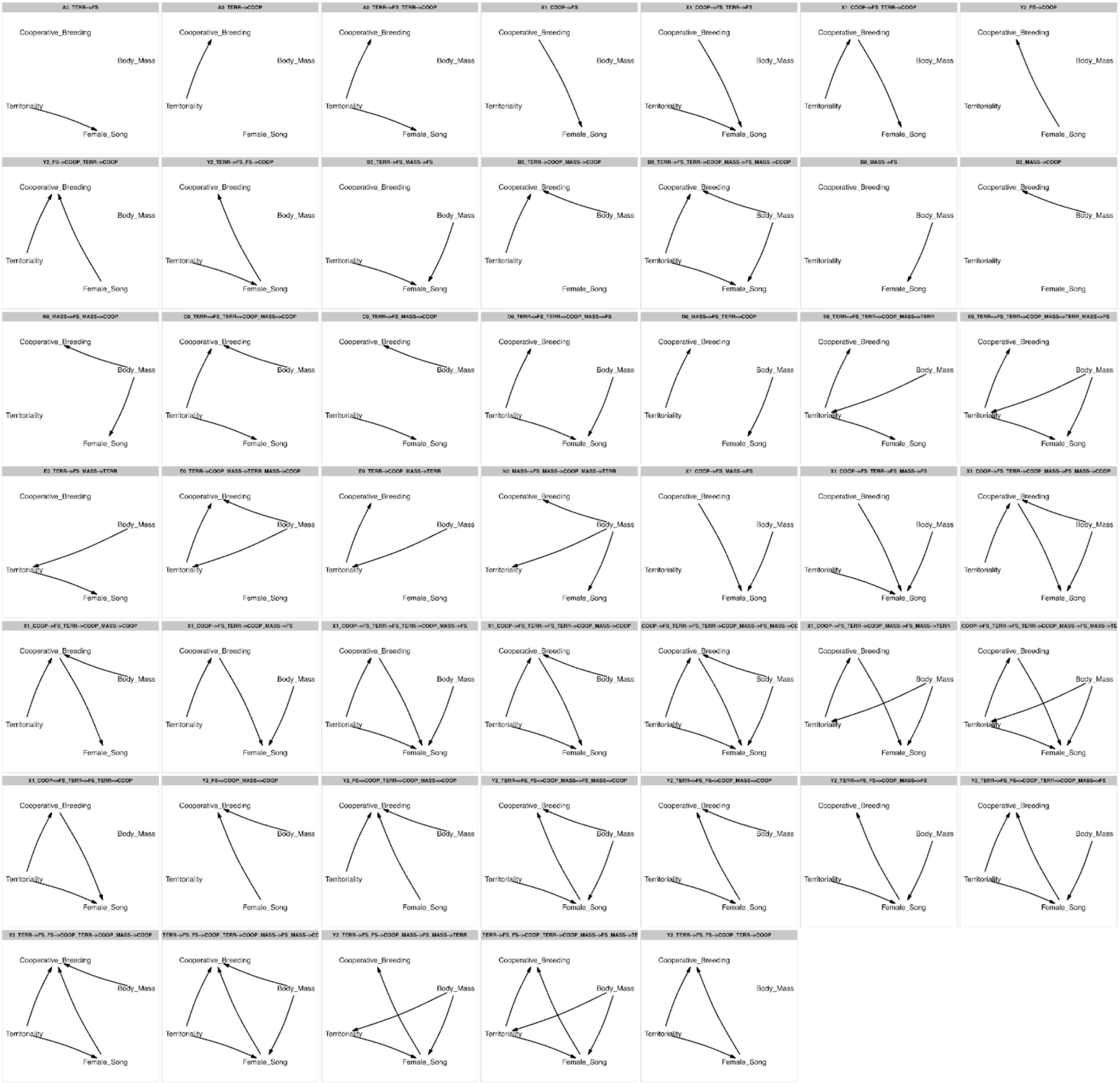

**Supplemental Figure 2:** Transition plots comparing evolution of female song with other sociality metrics: A) social group size larger than approximately 30 individuals, B) group size larger than pair, C) social bonds lasting longer than one year, D) social bonds lasting one season or longer, E) familial living, F) more than two caretakers, G) two or more caretakers, H) colonial living, and I) social mating system. Arrow weights vary as in Figures 4 and 5 and show the median log-scaled relative number of transitions that were estimated to occur across 500 simulations. Arrow color denotes the median difference from the expected number of transitions across simulations, with red indicating that the observed number of state switches was greater than expected, and blue indicating fewer state switches than expected. Asterisks indicate that the percent of simulations where the number of transitions is greater (red arrows) or less (blue arrows) than the calculated expected number of transitions for the simulation in question in >95% of simulations (**) or 90-95% of simulations (*). Sample sizes: A) n = 255 species, B) n = 255 species, C) n = 270 species, D) n = 270 species, E) n = 468 species, F) n = 270 species, G) n = 270 species, H) n = 270 species, I) n = 323 species.

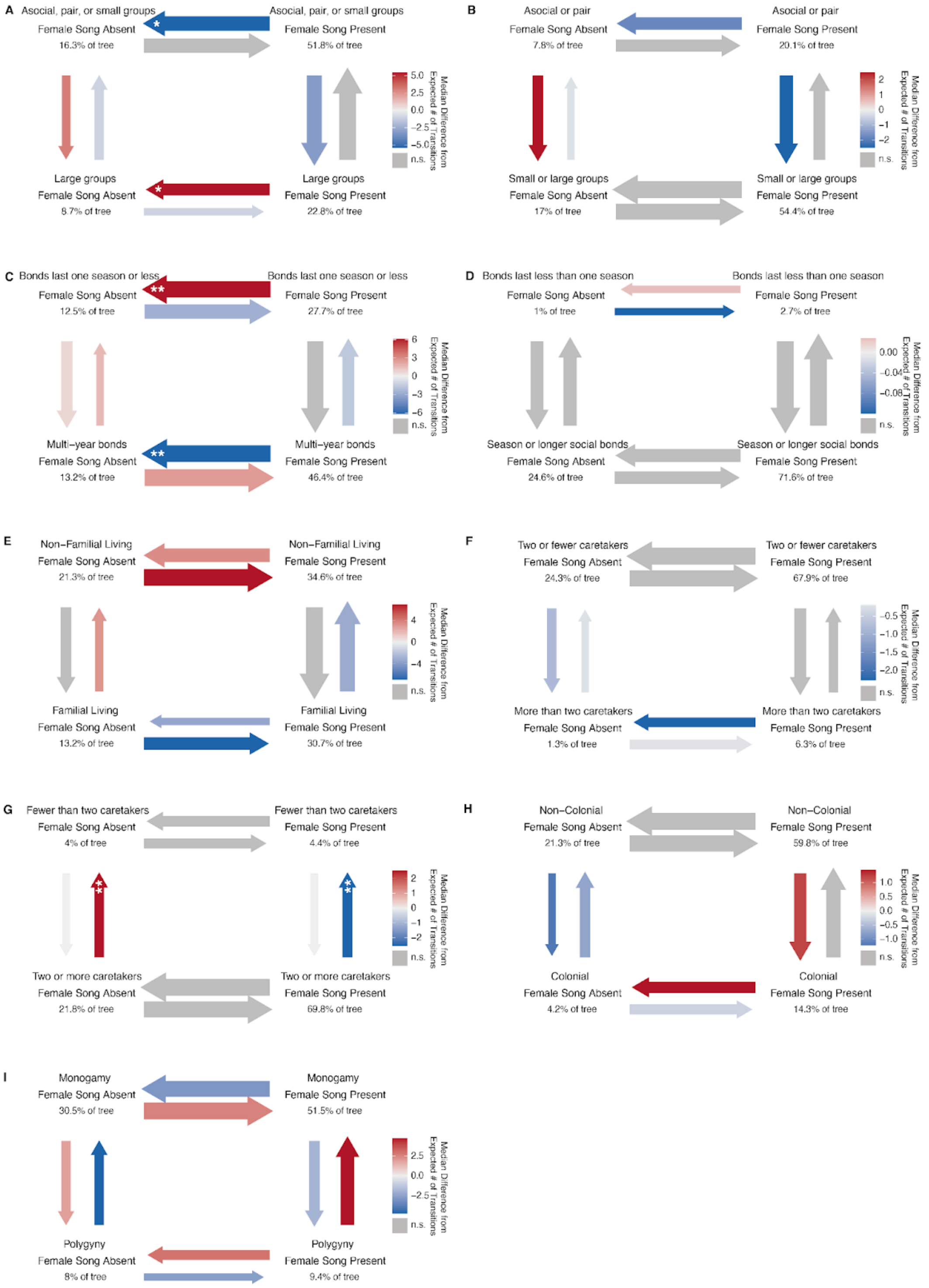

### Supplementary Results Text

*Data availability biases*

We found no evidence that cooperative breeding species are systematically more likely to have female song presence or absence data ($\chi^2 = 2.18$, $p = 0.14$; Table S13). Similarly, coloniality and polygyny showed no significant associations with either female song or cooperative breeding data availability (all $p > 0.05$). However, we detected significant biases related to territoriality, geography, and sexual dichromatism (Table S13). Territorial species were significantly more likely to have both female song and cooperative breeding data across all territoriality classifications (all $p < 0.001$). Holarctic species were overrepresented in female song databases ($\chi^2 = 56.8$, $p < 0.001$), and species with greater sexual dichromatism were more likely to have female song classifications ($t = -3.34$, $p < 0.001$). Importantly, among species with female song data, those classified as lacking female song had significantly higher sexual dichromatism ($t = 3.83$, $p < 0.001$), suggesting that dimorphism facilitates confident negative classifications rather than creating false positives for female song presence.

*Correlated evolution of female song and other social traits*

Several traits, though they did not significantly co-occur with female song in the binary trait co-occurrence analysis (Table ED1), nevertheless appeared to evolve non-independently with female song by affecting its likelihood of being lost or gained (Figure S2). We found that loss of female song was more likely to occur than expected in lineages with the largest social group sizes and that female song alongside a smaller group size is a relatively stable combination, with fewer transitions out of the state than expected (Figure S2A). Conversely, we found that long social bonds—those lasting longer than one breeding season—tended to promote transition to and maintenance of female song, while shorter social bonds tended to facilitate the loss of female song (Figure S2CB). This is consistent with the trends we observed when analyzing breeding system, considering that longer social bonds also tended to co-occur with cooperative breeding (Table S10). Familial living, which is associated with both cooperative breeding and longer social bonds, has a different pattern. We found that familial living tended to reduce the likelihood of either gaining or losing female song, while those likelihoods are increased in lineages with non-familial living (Figure S2E). Using the subset of species with data on number of caretakers from [21] reinforced our finding that lineages that had helpers at the nest, i.e. more than two caretakers, were less likely to lose female song, and those lineages without female song were less likely to increase the typical number of caretakers (Figure S2F). Intriguingly, the presence of female song appeared to greatly reduce the likelihood that a lineage would transition to fewer than two caretakers, whereas the loss of biparental care was more likely in the absence of female song (Figure S2G). There were not consistent evolutionary transition-rate trends between female song and colonial living or social polygyny (Figures S2H, S2I), although there is an interesting, albeit weak, trend suggesting that social monogamy with female song present as well as social polygyny with female song absent are both attractor states, i.e. that transitions into those states occur more frequently whereas transitions out of those states appear to occur less frequently.

*Accounting for other life history predictors*

We performed expanded phylogenetic generalized linear model analyses by iteratively adding other life history factors, including geographic region (holarctic vs. tropical; detailed sample sizes provided in Table S25), migratory behavior (non-migratory, partial migrant, migratory), familial living (familial vs. non-familial), plumage dichromatism (from sex-specific plumage scores), wing length dimorphism (from sex-specific morphometric data), and latitude (absolute value of the range centroid latitude) to the base models that were best supported for both effects on female song (best model: Female Song ~ Cooperative Breeding * Territoriality + Mass) and effects on cooperative breeding (Cooperative Breeding ~ Female Song * Territoriality). When we performed the stepwise iterative phylogenetic generalized linear model expansion analysis, we found that wing dimorphism and plumage dichromatism improved the model predicting female song presence by >2 AIC, while only familial living substantially improved the model predicting cooperative breeding (Tables S21,S22). In the expanded models, sexual dimorphism measures (wing: OR = 0.83, plumage: OR = 0.85) were negative predictors of female song presence. Even when including these additional predictors, the cooperative breeding-female song association remained robust (expanded FS model: CB OR = 5.78; expanded CB model: FS OR = 11.74; Tables S23,S24). In the expanded models with year-round territoriality, geographic factors emerged as significant predictors of female song: tropical species were less likely to have female song presence (OR = 0.67), while species at higher absolute latitudes were more likely to have female song presence (OR = 1.23). Familial living remained a strong predictor of cooperative breeding in both territorial classifications (OR = 10.62

with year-round territory vs. OR = 14.61 with weak/strong territory).
